# Defining the Layers of a Sensory Cilium with STORM and Cryo-Electron Nanoscopies

**DOI:** 10.1101/198655

**Authors:** Michael A. Robichaux, Valencia L. Potter, Zhixian Zhang, Feng He, Michael F. Schmid, Theodore G. Wensel

**Affiliations:** Verna and Marrs McLean Department of Biochemistry and Molecular Biology Baylor College of Medicine, Houston, TX, 77030; Graduate Program in Developmental Biology Baylor College of Medicine, Houston, TX, 77030; Medical Scientist Training Program, Baylor College of Medicine, Houston, TX, 77030; Current address: SLAC Linear Accelerator Laboratory, Menlo Park, California 94025

**Keywords:** Connecting cilium, rod photoreceptor, STORM, super resolution, Bardet-Biedl syndrome

## Abstract

Primary cilia are cylindrical organelles extending from the surface of most animal cells that have been implicated in a host of signaling and sensory functions. Genetic defects in their component molecules, known as “ciliopathies” give rise to devastating symptoms, ranging from defective development, to kidney disease, to progressive blindness. The detailed structures of these organelles and the true functions of proteins encoded by ciliopathy genes are poorly understood because of the small size of cilia and the limitations of conventional microscopic techniques. We describe the combination of cryo-electron tomography, enhanced by sub-tomogram averaging, with super-resolution stochastic reconstruction microscopy (STORM) to define substructures and subdomains within the light-sensing rod sensory cilium of the mammalian retina. Longitudinal and radial domains are demarcated by structural features such as the axoneme and its connections to the ciliary membrane, and are correlated with molecular markers of these compartments, including Ca^2+^-binding protein centrin-2 in the lumen of the axoneme, acetylated tubulin forming the axoneme, the glycocalyx extending outward from the surface of the plasma membrane, and molecular residents of the space between axoneme and ciliary membrane, including Arl13B, intraflagellar transport proteins, BBS5, and syntaxin-3. Within this framework we document that deficiencies in the ciliopathy proteins BBS2, BBS7 and BBS9 lead to inappropriate accumulation of proteins in rod outer segments while largely preserving their sub-domain localization within the connecting cilium region, but alter the distribution of syntaxin-3 clusters.

## INTRODUCTION

Primary cilia are thin membrane protrusions, each containing an axoneme with a 9+0 arrangement of microtubule doublets that arises from a basal body (BB) composed of a mother and daughter centriole located near the cell surface. They have been intensively studied in recent years because of their importance in sensory signaling, development, and physiological homeostasis. A host of inherited defects in cilium or basal body structure and function cause ciliopathies, a group of human diseases whose clinical symptoms include mental retardation, polydactyly, obesity, cognitive defects, cystic kidneys and blinding retinal degeneration (Badano et al., 2006; Ware et al., 2011).

Proteomic studies of cilia (Liu et al., 2007a; Zhao et al., 2016) along with genetic studies in humans and model organisms (Dean et al., 2016; Gherman et al., 2006; Ishikawa et al., 2012; Pazour, 2004) have revealed that mammalian primary cilia contain hundreds of different polypeptide components. These studies have provided some hints at the functions of cilia proteins, but in most cases their precise locations and functions are not known. These proteins include molecules necessary for precisely regulated transport of molecules into and out of cilia, such as the IFT proteins of the intraflagellar transport complexes, as well as numerous receptors, ion channels and transporters, and structural proteins.

Photoreceptor cells in the vertebrate retina contain highly specialized sensory cilia known as outer segments (OS) (Pearring et al., 2013; Wensel et al., 2016), which are cylindrical structures roughly 20 μm in length and 1.5 μm in diameter, packed with membrane disks containing the proteins of the phototransduction cascade (Wensel, 2008). They are connected to the inner segment (IS) of the cell by a short (~ 1.1 μm)**4** structure known as the connecting cilium (CC), which contains an axoneme extending from a basal body (BB) complex in the inner segment (IS) of the cell, and resembles the transition zone of other cilia (Pazour et al., 2002). Structurally, the microtubule doublets of the axoneme within the CC are surrounded by a 50 nm layer of cytoplasm and enclose a 150 nm central lumen for a total CC diameter of ~300 nm (Besharse et al., 1985; Gilliam et al., 2012). The requirement for daily renewal of ~10% of the membrane mass in the OS through transport of material through the CC (Wang and Deretic, 2014; Young and Droz, 1968) and the high metabolic demands of the phototransduction apparatus place particularly stringent demands upon this structure, as evidenced by the frequency with which ciliopathies include retinal degeneration among their symptoms (Bujakowska et al., 2017; Estrada-Cuzcano et al., 2012). Ciliopathic retinal degeneration follows a general time course of photoreceptor cilia dysfunction, followed by progressive rod cell death and blindness (Adams et al., 2007). Understanding the structure and organization of the rod sensory cilium and particularly the CC is thus important in order to understand these blinding diseases. These structures also serve as highly informative models for mammalian primary cilia, as they are particularly accessible to studies of structure, function, and genetics.

Two groups of supramolecular complexes which are important for photoreceptor function and health and also implicated in conserved transport mechanisms among eukaryotic cilia are the intraflagellar transport, IFT, complexes, and the BBSome complex formed by proteins associated with Bardet-Biedl Syndrome (BBS). BBS is a retinal ciliopathy caused by mutations to over a dozen BBS genes. The BBSome is a large protein complex composed of eight BBS ciliary proteins: BBS1, BBS2, BBS4, BBS5, BBS7, BBS8, BBS9, and BBIP10 (Loktev et al., 2008; Nachury et al., 2007), that are chaperoned by other essential BBS proteins including BBS17/LZTFL1 (Leucine-Zipper Transcription Factor Like 1) (Seo et al., 2011). Bbs mutant mouse lines universally model ciliopathic defects, including progressive retinal degeneration, which leads to near complete loss of the photoreceptor layer by 6 months of age (Davis et al., 2007; Mykytyn et al., 2004; Zhang et al., 2013). Rod cell death in these models has been attributed to gradual apoptosis caused by early defects in BBSome trafficking and subcellular morphological defects. *Bbs4*^-/-^ mutant mice have TUNEL-positive rods in the retina by 6 weeks of age (Mykytyn et al., 2004). At this same early stage, 35 nm vesicles are accumulated near the base and backed up into the lumen of the CC in *Bbs4*^-/-^ mutant rods (Gilliam et al., 2012), while in *Bbs1*^*M390R*^ and *Lztf1l/Bbs17*^-/-^ mutant models, significant accumulations of membranes were observed in the OS near the distal end of the CC (Datta et al., 2015).

The two complexes of IFT proteins (IFT-A and IFT-B), are prominent transport proteins in primary cilia, and form large “trains” that bind to microtubule motor proteins kinesin-II and dynein to actively traverse the microtubules of the axoneme and deliver and retrieve cargo proteins to the primary cilia tip (Nachury, 2014). These complexes are essential in rod cilia, as *Ift88/Tg737* mutant mice also have severe rod ciliopathic defects and retinal degeneration (Pazour et al., 2002). In primary cilia model live imaging experiments, the BBSome was determined to traffic along the axoneme at the same rate bidirectionally as IFT proteins (Lechtreck et al., 2009; Liew et al., 2014; Nachury et al., 2007; Ou et al., 2005), implying a BBSome-IFT interaction in primary cilia and rod cilia, and possible shared trafficking mechanisms for these proteins, which are disrupted in ciliopathies.

The precise functions of the BBSome and other complexes formed by ciliopathy proteins are not known, and even their precise locations within the rod cilium are not well defined. A major limitation has been the lack of an accurate map of the nanoscale molecular organization of the CC and surrounding regions. Immunofluorescence has been used to determine locations of numerous ciliopathy proteins, but only with rather low resolution because the entire 300 nm width of the CC covers little more than one Airy disk for a diffraction-limited spot in a confocal instrument. Conventional electron microscopy has provided a somewhat fuzzy picture of the ultrastructure of the CC (Wensel et al., 2016), confounded by the distortions introduced by fixation, embedding and ultra-thin sectioning, and the ambiguity imposed by multiple contrast images in samples stained with heavy metal salts.

Two complementary nanoscopy techniques that overcome many of these limitations are cryo-electron tomography and super-resolution single-molecule localization fluorescence. Cryo-electron tomography has provided important new insights into the structure of the rod cell and its connecting cilium (Gilliam et al., 2012; Nickell et al., 2007), but previous studies have not made use of the dramatic improvements in signal-to-noise and resolution that can be obtained through the technique of sub-tomogram averaging (Schmid, 2011). This technique has been applied with great success to eukaryotic flagella (Bui and Ishikawa, 2013; Koyfman et al., 2011; Nicastro et al., 2011), which are motile cilia, and to centrioles (Li et al., 2012) but is just beginning to be applied to mammalian primary cilia (Wensel et al., 2016).

Advances in super-resolution fluorescence nanoscopy have made it possible to localize specific proteins at resolutions well below the diffraction limit. Stochastic optical reconstruction microscopy (STORM) is a photoswitching nanoscopy in which samples are immunolabeled with antibodies conjugated to organic “photoswitching” fluorophores that cycle from a dark state to an active emitting state, and switch off again, repetitively. STORM experiments capture these single molecule photoswitching events randomly over the course of acquisition period, after which the point spread function (PSF) from each of these events are reconstructed as individual Gaussian data point localizations that are collected into reconstruction maps with 20-30 nm resolution in the X-Y dimension (Bates et al., 2007; Rust et al., 2006).

In this work we have used cryo-electron tomography and sub-tomogram averaging to define the three-dimensional architecture of the basal connecting cilium and centriole. We then combined the results with STORM imaging to define the radial subcompartments of the CC via sub-diffraction level reconstruction of immunotargets. Using this approach, the molecular constituents of the CC were reconstructed into 4 distinct layers of the CC, and the effects on their organization and localization of BBS mutations were determined.

## RESULTS

### Structure of Connecting Cilium by **Cryo-ET and Subtomogram Averaging.**

In order to determine the repeating structural features of the CC and define the subdomains’ geometries, we collected cryo-ET data on rods as described previously (Gilliam et al., 2012; Wensel and Gilliam, 2015; Wensel et al., 2016), and selected an unflattened region of the basal one-third of a CC for subtomogram averaging, based on nine-fold symmetry, using an approach previously used to refine the structure of a mouse rod daughter centriole (Wensel et al., 2016). The resulting map (Figure 1) showed good signal-to-noise and many clearly resolved nanometer scale features from the base of the mother centriole through the entire length of the selected region of the axoneme. These include the microtubule triplets of the centriole, the terminal plate in the lumen of the axoneme, the microtubule doublets of the axoneme, the ciliary membrane, and connections between the microtubules and the membrane. There are numerous filamentous and planar structures emanating from the mother centriole of the basal body, which are reinforced by 9-fold averaging; these bear little resemblance to the pinwheel-like or weather-vane-like structures referred to as “distal appendages,” “transition fibers” or “mylar sheets” in conventional TEM images of other basal bodies, but may be similar in function and molecular composition. There are also connections between the basal region of the axoneme and the ciliary membrane that display ninefold symmetry, and likely correspond to the structures commonly referred to as “Y-shaped links” or “champagne glass structures.” (Figure 1G, 1H) Although these do have a Y-shaped appearance in certain projection views, they actually consist of two different types of connections, each type emerging from the axoneme at a different angle, and appearing at different axial positions. When viewed in three dimensions, neither of them can be accurately described as “Y-shaped.”

**Figure 1.**
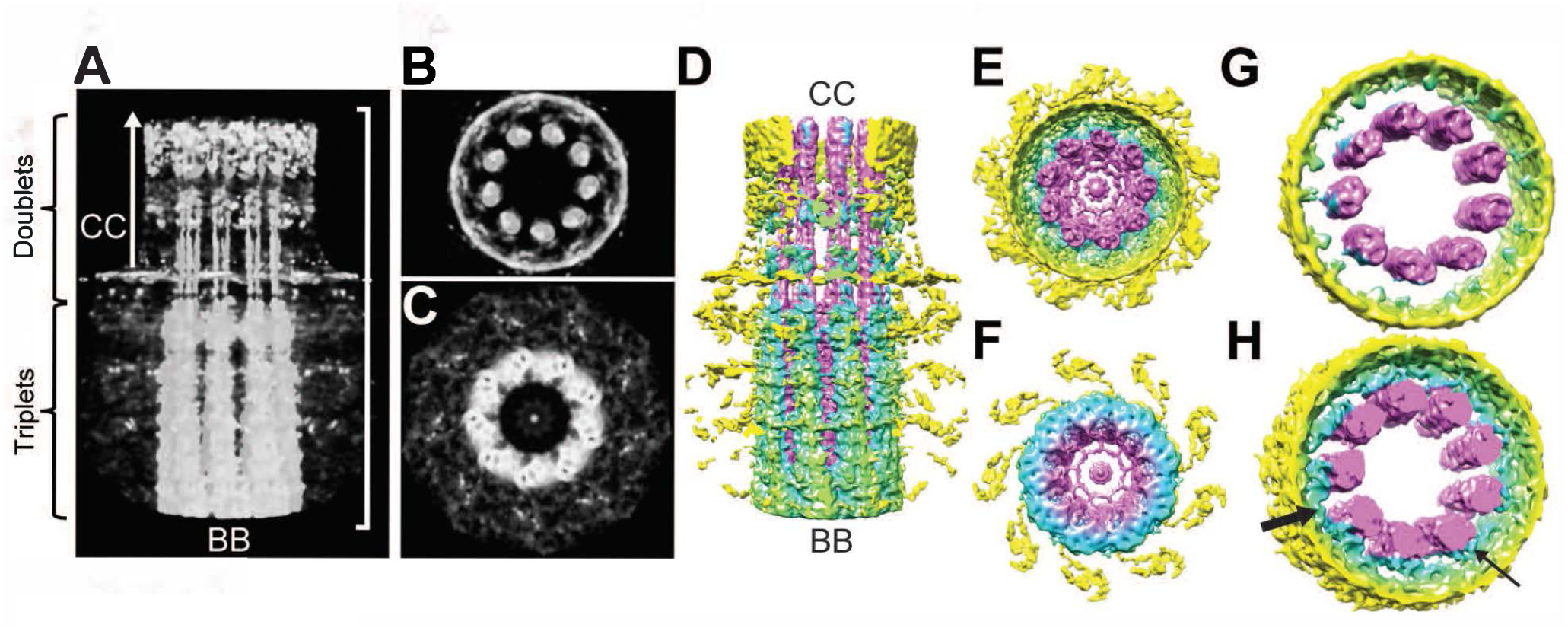
Structure of connecting cilium and basal body by cryo-ET and subtomogram averaging. (A-C) Volume-rendered representations of the final map after 9-fold averaging and 15 nm low-pass filtering. Vertical scale bar on right side of panel A is 700 nm and applies to panels A-F. (D-F) Projection views of entire map. Surface representations of entire map from the side (D), top (CC end, E), or bottom (BB end, F). Threshold for F was set slightly higher than for F to visualize the lumens of the microtubules. The terminal plate, and filamentous/planar structures emanating from the BB are observed within the lumen. (G), (H) Surface representations of basal portion of CC showing the presence of two distinct types of connections between the axoneme and plasma membrane at lower (thick arrow) and more distal (thin arrow) axial positions in panel H. Surface representation in panel G is the portion immediately above the one shown in panel H, revealing the absence of visible connections between the axoneme and plasma membrane in the symmetrized map.

This map defines several clearly defined structural domains. These include the mother centriole, containing triplet microtubules and a lumen, along with protruding appendages; the axoneme composed of microtubule doublets, the lumen of the axoneme, the volume between the axoneme and the ciliary membrane, the membrane itself, and the glycocalyx extending from it. This last feature is not visible in our map, likely due to its lack of nine-fold symmetry and relatively low electron density. These clearly demarcated structural domains provide a framework for interpreting immunolocalization results in the context of a three-dimensional model.

### Super-resolution Fluorescence Nanoscopy of Rod Cilium Domain Markers

In order to define subdomains in molecular terms, we tried several approaches to acquiring STORM data from immunostained mouse rods, beginning with a preparation used previously to determine structural features of the rod cilium by cryo-electron tomography (Gilliam et al., 2012; Wensel and Gilliam, 2015), and including methods we used previously for immunolocalization in retinal whole-mounts (Gilliam et al., 2012; Wensel and Gilliam, 2015) or cryo-sections (Gilliam and Wensel, 2011; He et al., 2016; Mojumder et al., 2009), with a variety of fixation and membrane permeabilization methods. These all suffered from either poor photon yield or poorly preserved cellular morphology. We eventually developed a protocol adapted from one used previously to image retinal interneurons (Sigal et al., 2015) in which retinas are labelled with organic-dye conjugated antibodies, followed by embedding and sectioning with an ultramicrotome, and etching with sodium ethoxide. This procedure yields good photoswitching with reasonable photon yields and good preservation of cell morphology. The imaging optics and STORM localization analysis procedure generated two dimensional reconstructions with single molecule resolution on the order of 20 nm in x and y, and a depth of field that yields a projection through as much as an entire CC in the z dimension (~300 nm, Figure 2).

**Figure 2.**
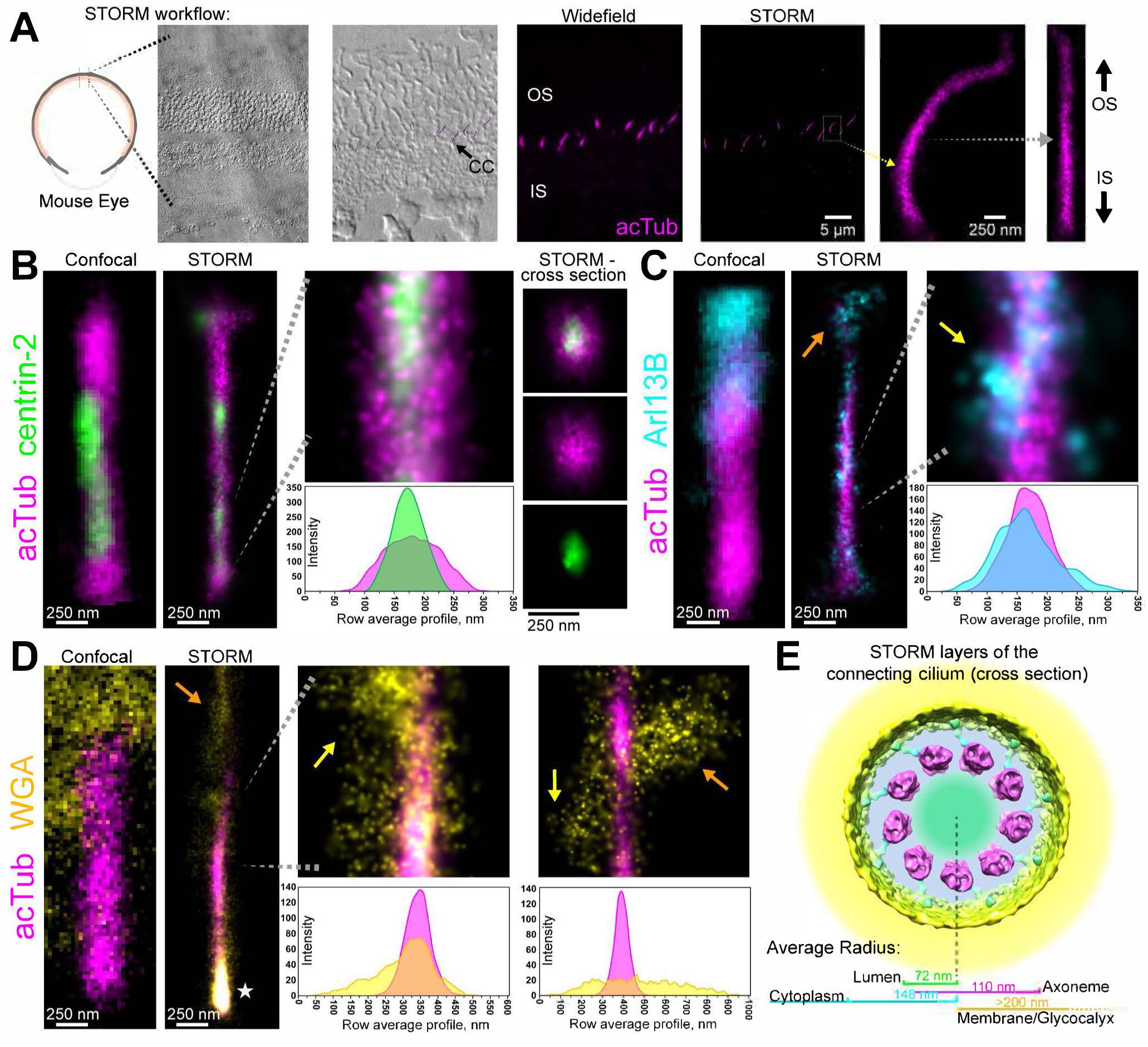
STORM reconstruction of 4 distinct layers of the rod connecting cilium. (A) Workflow of STORM experiments in mouse retina. A differential interference contrast (DIC) image of the retina is magnified and overlaid with a reconstruction of acetylated-alpha tubulin (acTub) to illustrate the location of the connecting cilia (CC) clusters in the tissue section. This same STORM reconstruction is compared to a widefield image, and a CC acTub cluster from the STORM reconstruction is isolated (yellow dotted arrow) and digitally straightened (gray dotted arrow). All CC clusters shown are oriented with the top of the CC directed toward the outer segment (OS). (B–D) Reconstructions of clusters for sub-domain markers in 2–color STORM experiments: immunostaining for acetylated α-tubulin (acTub, magenta, B–D) to mark the axoneme; for centrin-2 (green, B) to mark the lumen; for Arl13B (cyan, C) to mark the ciliary cytoplasm; and wheat germ agglutinin (WGA, yellow, D) to mark the ciliary membrane and glycocalyx. Confocal images of CC that are immunostained with the same antibodies as adjacent STORM reconstructions are depicted at the same scale for comparison. Row average profiles are arranged directly below their corresponding STORM reconstruction images. (E) Diagram overlaying STORM compartment localization results in a cross section view of the CC with a cryo-ET cross-section structure to scale, as well as the average radial measurements from STORM images for each CC layer. Yellow arrows = STORM clusters in CC. Orange arrows = STORM clusters in OS. Stars = basal body/pericentriolar region.

To calibrate our localization accuracy and validate the resolution, we used two markers whose localization is well characterized within the cilium: acetylated alpha-tubulin (acTub), which is a component of the microtubule doublets on the axoneme, and centrin-2, a calcium binding protein known to occupy the lumen of the axoneme (Giessl et al., 2006; Pulvermuller et al., 2002; Trojan et al., 2008) (Figure 2B).

Based on the inside diameter of the axoneme determined by cryo-electron tomography to be 156 nm, and a nominal STORM resolution of 25 nm, we might expect the profile of the luminal centrin-2 staining to have a profile with a width of approximately 181 nm. Surprisingly, the observed width (distance between profile positions with amplitudes of 1/e times the maximum) is only 143 nm ± 17 nm (Figure 2B, S1, n=11). This result leads to two striking conclusions: 1) that localization can indeed be determined with resolutions of tens of nm, and 2) centrin-2 is confined to the central region of the axoneme, and does not extend to the microtubule walls. This narrow range suggests that centrin-2 may be a component of the proteinaceous particles previously observed within the central lumen of the axoneme by cryo-electron tomography (Gilliam et al., 2012). Not surprisingly, acTub signal has a wider profile, consistent with it forming the microtubules surrounding the lumen, and the corresponding width is 218 nm +/- 32 nm (from all experiments in Figure 1, S2, S3, n = 44), consistent with a structure 200 nm wide, labelled with both a primary and secondary antibody that could extend beyond the surface of the microtubules by as much as 20 nm on either side. As with centrin-2, the profile confirms the super-resolution of our imaging methods. In contrast to the expectation for signal arising strictly from the axonemal microtubules, the profile of acTub often displays a peak, rather than a trough in the center of the axoneme, suggesting the possible presence of a pool of soluble acetylated tubulin in the lumen. Indeed, in cross-sectional views (e.g., far right panel of Figure 2B), the central location of the centrin-2 within the acTub staining is apparent, but the acTub signal appears uniform across the lumen. In select portions of the CC, however, the acTub profile is more consistent with signal coming predominantly from the microtubules (Figure S1B).

The small GTPase, Arl13B, a cilium protein with a less well established sub-ciliary localization (Bhogaraju et al., 2013; Cantagrel et al., 2008; Hori et al., 2008; Liem et al., 2012) served as a marker for the ciliary membrane and the cytoplasmic domain outside the axoneme (Figure 2C, S1C). The signal from this antigen was much less uniform, and extended well beyond the edge of the axoneme, consistent with association of this palmitoylated protein (Cevik et al., 2010; Roy et al., 2017) with the ciliary membrane. In contrast to its pattern in some epithelial cells (Cevik et al., 2013), Arl13B is not restricted to a subdomain of the CC, but is found throughout its length. However, unlike the centrin-2 and acTub labeling, and in contrast to its appearance at confocal resolution (*e.g*., left-most panel of Figure 2C), the signal was not distributed uniformly along the length of the CC, and appeared primarily in clusters at irregular spacings along the long axis. This clustering has important implications for Arl13B function, suggesting its association with large sparse complexes along the ciliary axis. It has been reported to act as a guanine nucleotide exchange protein for a related small GTPase, Arl3 (Gotthardt et al., 2015; Humbert et al., 2012; Ivanova et al., 2017), which in turn plays a critical role in releasing farnesylated proteins from complexes with PDE6δ/prenyl-binding protein during ciliary trafficking. In addition to the consistently strong labeling of the CC, Arl13B signal was also seen in the proximal outer segments, with a diffuse pattern not confined to the region near the axoneme (orange arrows in Figures 2C, S1C).

To assess the signal for a marker that should be external to the ciliary membrane, we used the lectin wheat germ agglutinin (WGA) to label the glycocalyx of the cilium. In previous immuno-EM studies, WGA-gold conjugates strongly labeled an irregular glycoconjugate cloud that partially enveloped the CC of rods (Besharse et al., 1985; Horst et al., 1987). The STORM reconstruction of WGA was more than 700 nm wide in some cases, consistent with a glycocalyx extending hundreds of nm beyond the ciliary membrane (Figure 2D, S1D).

### Localization of IFT Complexes

Next, IFT81 and IFT88 proteins were targeted as subunits of the highly conserved core of the IFT-B primary cilia complex. With conventional fluorescence microscopy in rods, IFT protein signal is generally concentrated at either end of the CC with less clear localization along the CC axoneme (Liew et al., 2014; Pazour et al., 2002), whereas IFT88 subcellular localization via immuno-EM was prominently along the outer face of the axoneme microtubules (Sedmak and Wolfrum, 2010). Notably, IFT81 is a component of an IFT tubulin-binding module, which may establish the near colocalization of the IFT complex with the axoneme (Bhogaraju et al., 2013). Both kinesin-II, the anterograde motor protein for the IFT complex, and dynein, the retrograde motor, have been similarly immunolocalized to the CC (Beech et al., 1996; Mikami et al., 2002). We observed a very bright signal around the BB for IFT81 and IFT88, as well as along the axoneme in the OS. Multiple IFT clusters were observed along the CC (Figure 3A, 3B, S2), and these STORM clusters were somewhat larger and less numerous than the clusters observed for Arl13b, but were located within the same radial domain. Averaged along the length of the CC, the average radii were 143 nm ± 25 nm (IFT81, n=13) and 138 nm ± 30 nm (IFT88, n=20). These radial localizations are nearly identical to those observed with Arl13B (148 nm ± 24 nm, n=8), which is consistent with a reported in vitro Arl13b-IFT-B complex interaction (Cevik et al., 2013). The large size of the IFT81 and IFT88 STORM clusters suggests they likely represent IFT particles, or “trains”, and in some cases these are very closely associated with the surface of the axoneme (Figure 3A); in others, they fill the entire radial span between the axoneme surface and the ciliary membrane (Figure 3B), consistent with immunoelectron microscopy results showing IFT staining of the axoneme and the ciliary membrane (Sedmak and Wolfrum, 2011).

**Figure 3.**
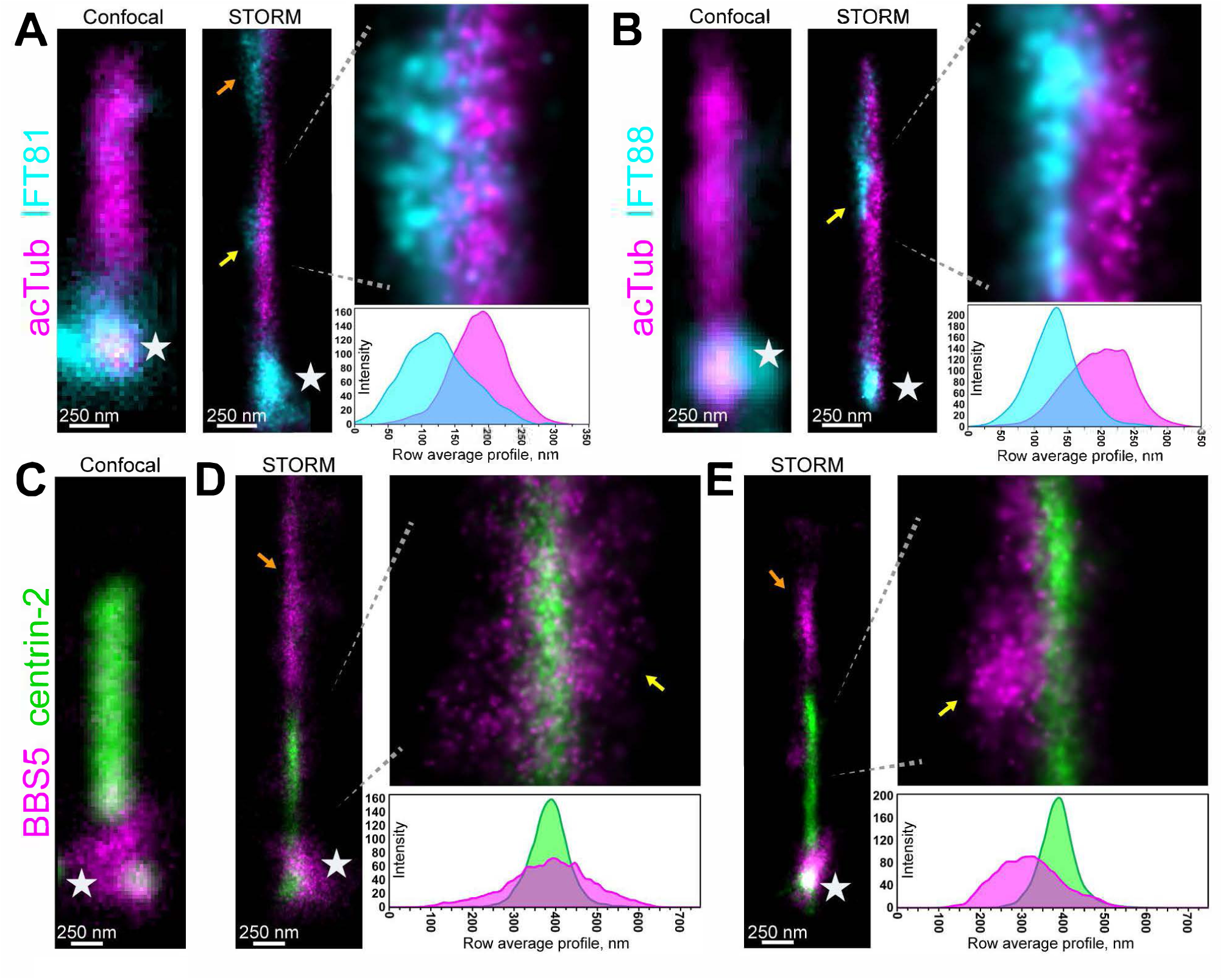
Super-resolution localization of IFT proteins and BBS5. (A) STORM reconstructions for acTub (magenta) and IFT81 (cyan). (B) STORM reconstructions for acTub (magenta) and IFT88 (cyan). (C-E) Two examples of STORM CC reconstructions of centrin-2 (green) and BBS5 (magenta). For each antigen reconstructed with STORM at the CC, a confocal image of an identically immunostained CC is shown at the same scale for comparison. Magnified regions are presented with corresponding row average profiles aligned directly beneath. Yellow arrows = STORM clusters in CC. Orange arrows = STORM clusters in OS. Stars = basal body/pericentriolar region.

### Localization of the BBSome

To localize the BBSome we relied on a monoclonal antibody specific for the pleckstrin-homology (PH)-domain containing BBSome subunit BBS5, whose localization within the CC is unresolved (Smith et al., 2013). Out of many antibodies for BBS subunits we tested, this was the one that provided the most consistently reliable results under our labeling and imaging conditions, although STORM reconstructions with a BBS9 antibody matched and overlapped with the BBS5 localization pattern at the CC (Figure S3). To correlate BBS5 signal with the defined radial domains, we co-labeled for centrin-2. As shown in Figures 3 and S3, in WT retina, BBS5 is located in three subcellular regions at different levels, with very intense staining at the basal body complex (white stars), somewhat less intense staining along the axoneme in the outer segment (orange arrows), and less intense but consistently detectable clusters in the CC volume outside the axoneme. Somewhat surprisingly, the signal from these CC clusters extends in many cases considerably more than 150 nm from the center of the axoneme marked by centrin-2 (mean BBS5 radius = 220 ± 30 nm, n=19), so that BBS5 extends beyond the nominal 300 nm width of the CC, even if the extra ~40 nm width introduced by the antibodies and the localization uncertainty is taken into account. These appear to represent localized protrusions of the membrane into which the BBSome is inserted. Local membrane distortions would be consistent with the observation that the BBSome is able to form a coat on the surface of phospholipid bilayers (Jin et al., 2010). In contrast to the IFT proteins which are very closely associated with the BB, BBS5 staining near the BB is more diffuse, consistent with its previously reported association with the pericentriolar material.

### Co-localization of the BBSome and IFT proteins

To determine the extent of co-localization of BBS5 and IFT proteins within the CC, it was necessary to include a third label for alpha-tubulin 1 (TUBA1) to mark the CC and distinguish these proteins within the CC from their distinct populations in the inner and outer segments. As our experimental setup precluded simultaneous STORM imaging of the third label, we superimposed widefield images of TUBA1 staining on two-color super-resolution reconstructions of BBS5 and IFT88 (Figure 4). The results reveal that there are also membrane protrusions at the CC containing IFT88, with some protrusions containing both IFT88 and BBS5, and others containing only one or the other. Thus, there are at least four distinct populations of BBS5: one associated with the centrioles and pericentriolar material, a second associated with the axoneme in the outer segment, a third within the CC co-localized with, and possibly participating in complexes with IFT particles containing IFT88, and a fourth present in the CC in regions not containing IFT88. This diversity of locations and distributions supports proposed roles for the BBSome in dynamic trafficking of cargo in cooperation with IFT particles and observed continual movement of BBSome subunits observed in live cell imaging.

**Figure 4.**
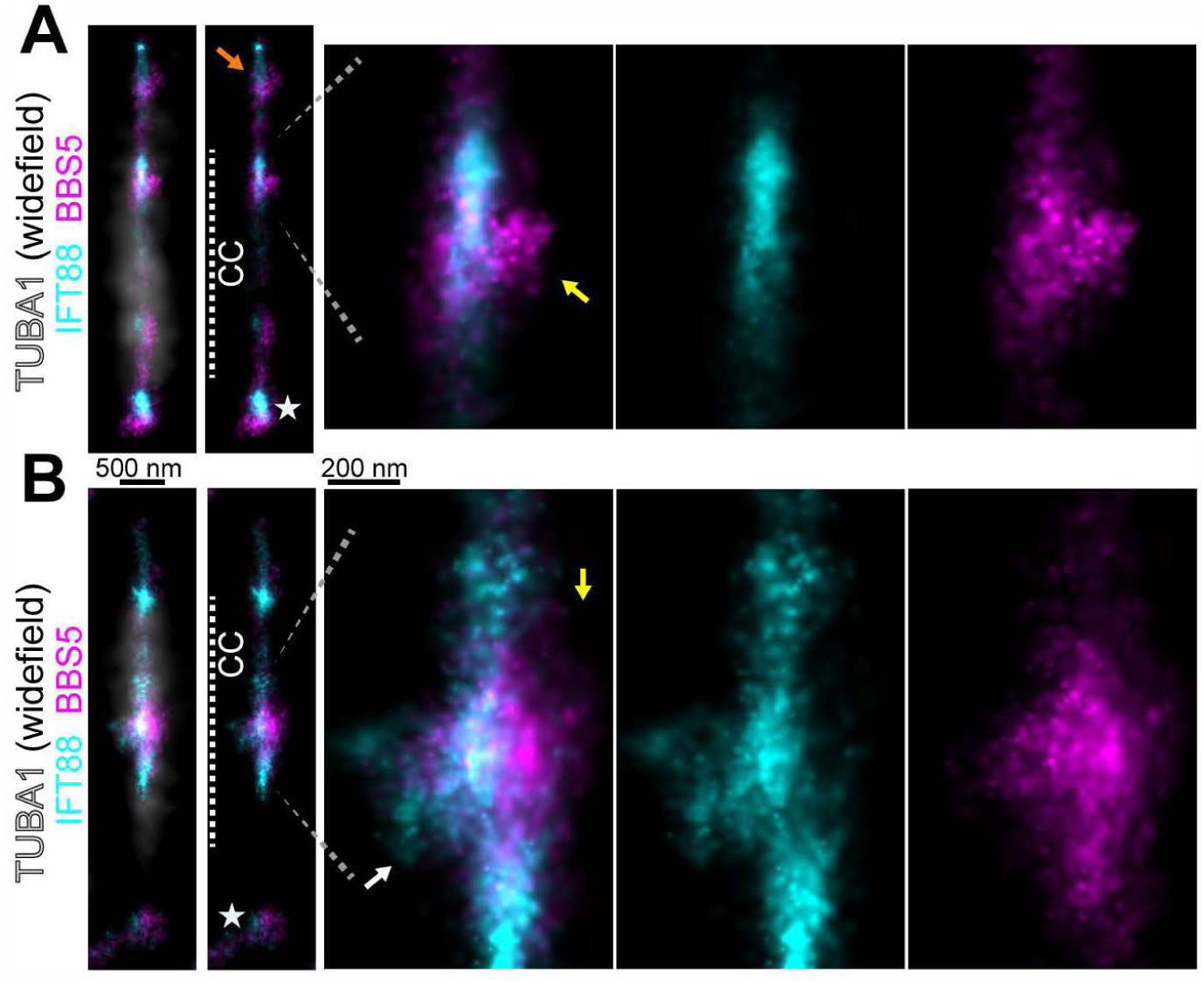
Colocalization of IFT88 and BBS5 along the connecting cilium with STORM. (A), (B) Two examples of isolated connecting cilia (CC) from overlaid 2-color reconstructions of IFT88 and BBS5 with a widefield fluorescence image of α-tubulin 1 (TUBA1) staining from the same region to mark the CC. See Figure S4 for full retinal view of this staining. The CC corresponding to the TUBA1 profile is marked in STORM-only images. Magnified regions of colocalized IFT88 and BBS5 clusters are from this region. Yellow arrows = BBS5 localization surrounding IFT88 clusters, White arrows = an IFT88 cluster isolated from BBS5, Orange arrow = colocalization of IFT88 and BBS5 along the axoneme within the OS, Stars = basal body/inner segment localization.

When BBS5 clusters were adjacent to IFT88 clusters, the were consistently patterned at the edge of IFT88 clusters and extended away from the CC as if surrounding the IFT88 clusters. This pattern is consistent with a model for transport machinery at the CC in which BBS5/the BBSome interacts with the IFT complex as part of a larger transport complex with a molecular arrangement of IFT proteins near the axoneme microtubules via motor proteins that are surrounded by BBSome complexes which extend out into the cytoplasm and to the CC membrane.

### Localization of Syntaxin-3

Syntaxin-3 (STX3) is a tSNARE transmembrane protein whose retinal expression is predominantly in the IS of rod cells (Figure S3D), including the region surrounding the BB at the base of the connecting cilium in *Xenopus* and rodent rods (Chuang et al., 2007; Mazelova et al., 2009). Furthermore, STX3 is proposed to be important for rhodopsin vesicle trafficking (Chuang et al., 2007; Mazelova et al., 2009), and is severely mislocalized in *Bbs* mutant rods (Datta et al., 2015) (see below, Figure S5B), which suggests that STX3 transport is dependent on a yet unknown function for BBS proteins and the BBSome in the CC.

In STORM images of the CC, clusters are observed for STX3 in three major locations (Figure 5, S3D): extensive signal at the BB (white stars) and pericentriolar region, large punctate clusters along the CC membrane, and some staining occasionally observed in the vicinity of the axoneme in the OS (orange arrows in Figure S3). The STX3 clusters at the CC are more compact than those observed for IFT proteins and BBS5, and are variable in size; as measured along the longitudinal axis of the CC the average length was 558 nm +/- 130 nm (n=14). Radially they extended as far as > 200 nm from the center of the CC. Thus, along the CC, STX3 was localized as large and cohesive STORM clusters that were located in variable and likely transient positions at the exterior CC membrane STORM layer, possibly as membrane transport structures in transit between the IS and OS.

**Figure 5.**
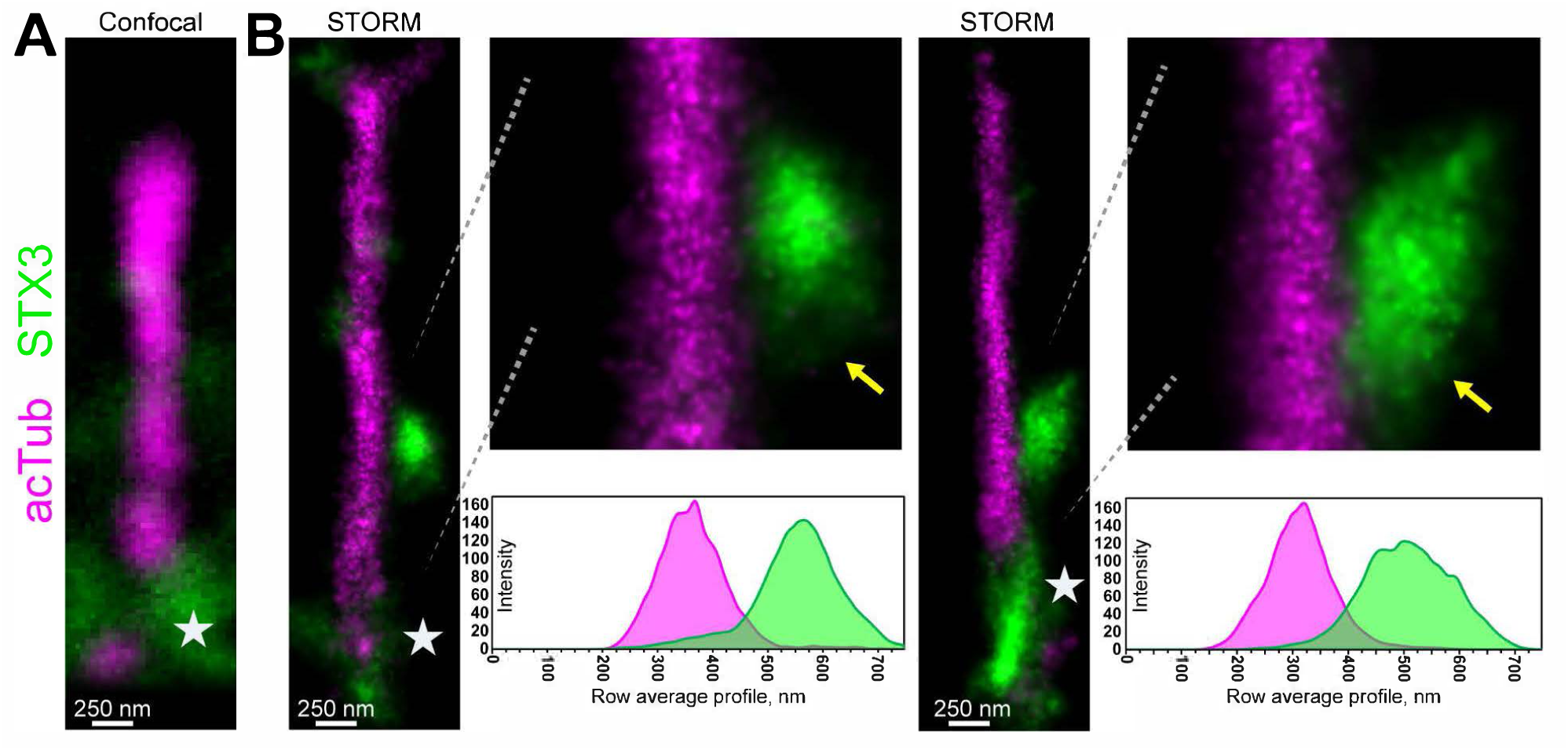
STORM reconstruction of syntaxin-3 (STX3) at the connecting cilia of wild-type mouse rod photoreceptors. (A) Confocal image of a connecting cilium (CC) immunostained for acetylated α-tubulin (acTub) and syntaxin-3 presented at the same scale for comparison. (B), (C) Two examples of STX3 STORM reconstructions at the CC. Magnified regions of the CC featuring large clusters STX3 are presented with corresponding row average profiles aligned directly beneath. Yellow arrows = Large STX3 STORM clusters localized along the CC. Stars = basal body/inner segment STX3 localization.

### Effects of *Bbs* gene Deficiencies on Localization

The eight BBS proteins of the BBSome (BBS1, BBS2, BBS4, BBS5, BBS7, BBS8, BBS9, and BBIP10), are expressed primarily as a full complex in rod sensory cilia as in ciliated mammalian cells; however, select BBSome subunits can form stable subcomplexes upon co-expression and during complex assembly (Nachury et al., 2007; Zhang et al., 2012), leading to the possibility of functional BBSome subcomplexes in rod cilia. Thus, we immunolabeled BBS5 for STORM reconstruction in rod cells from *Bbs2*^-/-^,*Bbs4*^-/-^ and *Bbs7*^-/-^ mutant mouse models of BBS retinal degeneration to determine if BBS5 and the remaining incomplete BBSome are mislocalized in these mutant backgrounds.

For these STORM experiments in mouse mutants with degenerating retinas, ages between 5 weeks and 8 weeks were selected as timepoints when enough rod cells remain intact for sufficient sampling size yet after sufficient aging to visualize maximal protein accumulation or mislocalization defects. *Bbs2*^-/-^ mutant mice have a rate of photoreceptor degeneration comparable to *Bbs4*^-/-^in which apoptosis was not apparent until 6 weeks; whereas, apoptosis was not measured in *Bbs7*^-/-^mice (Zhang et al.,2013). By 8 weeks, all *Bbs* mutant retinas demonstrated evidence of photoreceptor degeneration through retinal thinning; however, all mutant retinas still maintained numerous intact rod cells in the photoreceptor cell layer (Figure S4D).

BBS5 was not severely mislocalized to any particular photoreceptor region in any of the three *Bbs* mutant mouse strains compared to WT and heterozygous controls (Figure 4, 6, S5 & S6). In widefield fluorescence images, small abnormal BBS5 puncta were observed in the OS layer of the mutant retinas that were not present in wild-type and heterozygous controls (Figure S5A), and in STORM reconstructions some BBS5 mis-accumulation beyond normal axoneme localization is evident in the OS (Figure 6, S6). However, subciliary localization within the CC was unaffected in any of the *BBS* knockouts: BBS5 STORM localizations extended to the membrane STORM layer of the CC as in WT, and were found along the length of the CC. Surprisingly, BBS5 STORM clusters at the BB/IS were not significantly affected in any of the BBS mutant rods. These results indicate that BBS5 and BBSome subciliary localizations to the CC are maintained despite a degenerating background with known defects associated with *Bbs* ciliopathy models that precede impending rod cell death. Thus, incomplete BBSome complexes are at least partially functional and perhaps contribute to moderating defects in CC trafficking.

**Figure 6.**
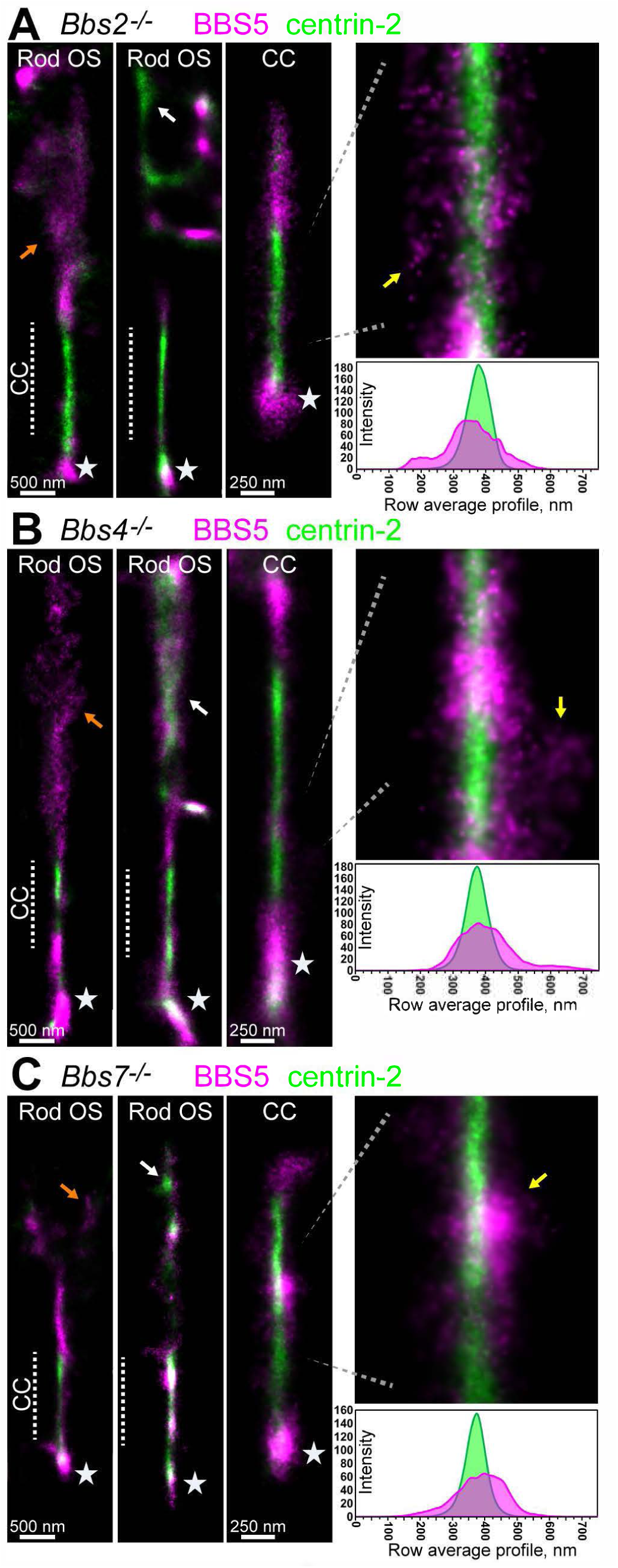
BBS5 and centrin-2 mislocalization in *Bbs* mutant mouse rods. Example STORM reconstructions for BBS5 (magenta) and centrin-2 (magenta) from (A) *Bbs2*^-/-^ rods (age = 8 weeks), (B) *Bbs4*^-/-^ rods (age = 8 weeks) & (C) *Bbs7*^-/-^ rods (age = 7 weeks). For each mutant, rod outer segment (“Rod OS”) STORM examples from a wider view are depicted to demonstrate BBS5 (orange arrows) and centrin-2 (white arrows) mislocalizations to the OS. In these reconstructions, the CC sub-region is indicated with a dashed line. Magnified STORM regions are presented with corresponding row average profiles aligned directly beneath. Yellow arrows = BBS5 STORM clusters along the CC. Stars = basal body/inner segment BBS5 clusters.

STX3 was previously found at aberrantly high levels in the OS of *Lzftl1/Bbs17*^-/-^knockout mice and grossly mislocalized to the OS layer of retina in *Bbs17*^-/-^ and *Bbs1*^*M390R*^ mutant mice (Datta et al., 2015). In addition, rod cells from these mutants featured large accumulations of membranes at the base of the OS near the distal end of the CC. Thus, rod cell localization of STX3 and regulation of its transport through the CC is regulated in some manner by the BBSome. Therefore, we used STORM to localize STX3 in rods from *Bbs2*^-/-^,*Bbs4*^-/-^ and *Bbs7*^-/-^ mutant mice to determine if any subciliary localization differences of STX3 in the CC are caused by disrupted BBSome composition and function.

In STORM reconstructions of STX3 from each *BBS* mutant line (Figure 7), STX3 was grossly mislocalized from the IS to the OS compared to WT or *Bbs* heterozygous controls (Figure S5B). This pattern of mislocalization is also apparent in STORM reconstructions of individual rod cells of all of the *Bbs* mutants, where STX3 clusters in the OS were typically punctate and located along the edge of the OS membrane as sometimes diffuse localized throughout the entire OS of mutant rods (Figure 7, S7). Despite OS accumulation, STX3 was normally localized to the BB in each *Bbs* mutant. Along the CC, the overall radial distribution of STX3 localization to the exterior membrane layer of the CC was maintained; however, *Bbs* mutant STX3 clusters were either more diffuse, less structured, or smaller than the large WT STX3 clusters (Figure 5).

**Figure 7.**
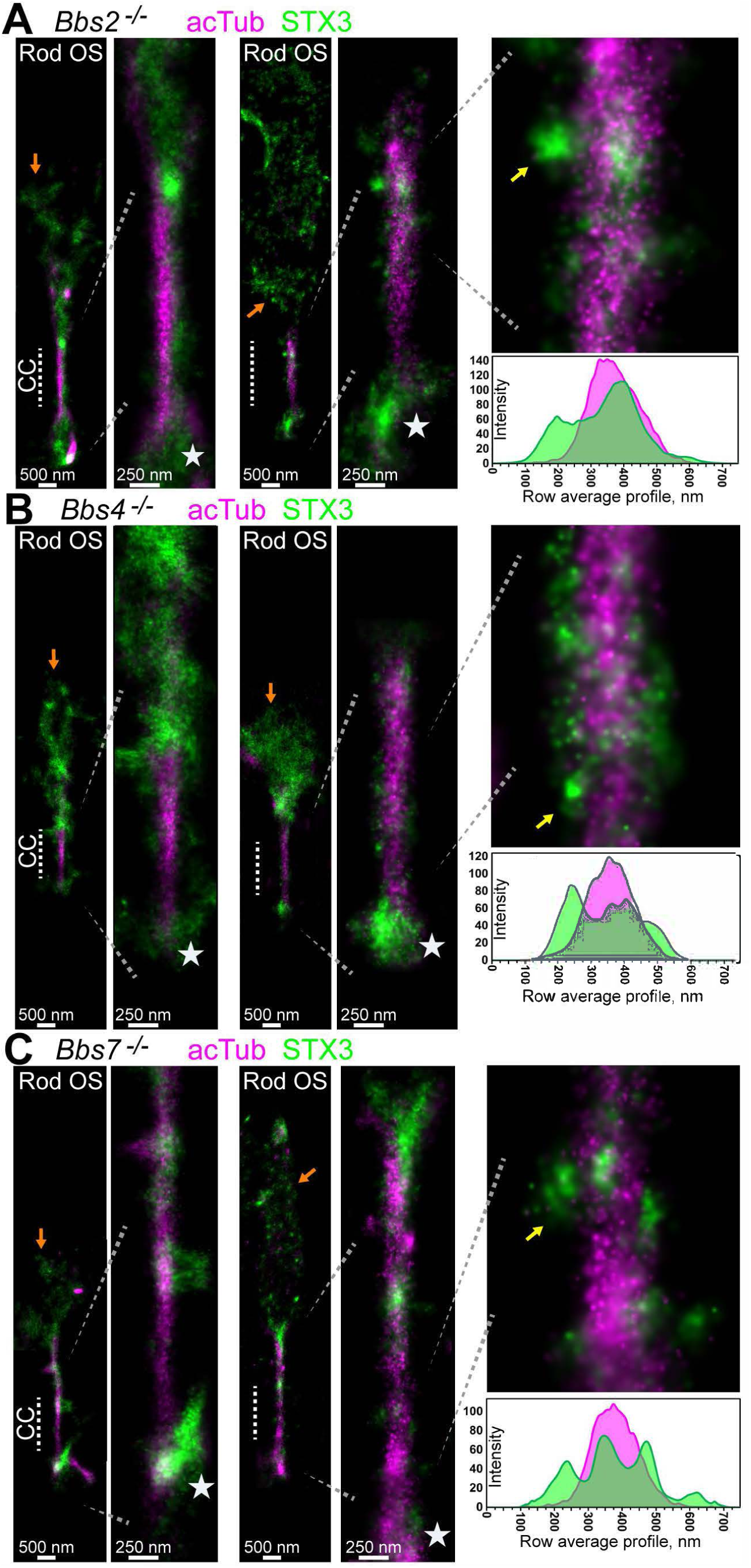
Mislocalization of syntaxin-3 in *BBS* mutant mouse rods. Example STORM reconstructions for syntaxin-3 (STX3, green) and acetylated α-tubulin (acTub, magenta) from (A) *Bbs2*^-/-^ rods (age = 8 weeks), (B) *Bbs4*^-/-^ rods (age = 5 weeks) and (C) *Bbs7*^-/-^ rods (age = 8 weeks). For each mutant, rod outer segment (“Rod OS”) STORM examples from a wider view are depicted to demonstrate syntaxin-3 (STX3) mislocalization to the OS (orange arrows). Magnified STORM regions are presented with corresponding row average profiles aligned directly beneath. Yellow arrows = STX3 STORM clusters along the CC. Stars = basal body/inner segment STX3 localization.

While the cluster organization of STX3 at the CC is clearly disrupted in Bbs mutant rods, radial localization of STX3 was only slightly altered in *Bbs7*^-/-^ cilia compared to WT (Fig. S5). Therefore, based on these STORM results in three different *Bbs* mutant mouse lines, subciliary localization of STX3 is only partially attenuated in the CC membrane layer despite gross OS accumulation. As such, anterograde and possibly partial retrograde trafficking evidently continue to some degree despite an incomplete BBSome composition and ciliopathy-related OS stress caused by protein accumulation and other mutant defects.

## DISCUSSION

Our application of nine-fold subtomogram averaging to examine the rod sensory cilium has yielded improved signal-to-noise for the symmetric features of the CC and the basal body complex, and revealed some previously unknown features. We previously applied this approach to a rod daughter centriole (Wensel et al., 2016), and the fundamental arrangement of the triplet microtubules is very similar in the two maps. Missing from both maps are triangular stain-accumulating structures extending from the basal body in a nine-fold pin-wheel arrangement as seen in many electron micrographs from basal bodies in other ciliated cell types or in immature mammalian rods (Greiner et al., 1981; Sedmak and Wolfrum, 2011). These are variously referred to as distal and subdistal appendages, “transition fibers” or “alar sheets.” There are structures extending from the mother centriole that have been reinforced by nine-fold symmetry, but these are irregular in appearance apart from their nine-fold pin-wheel arrangement. It is thus unclear what the relationship is, if any between the appendages seen in immature and mature basal bodies.

More well-defined structures are seen connecting the axoneme to the membrane in the basal region of the CC. In a previous cryo-ET study (Gilliam et al., 2012), numerous fibers or filaments were observed leading from the microtubules to the membrane, but these lack nine-fold symmetry and thus are not observed in the averaged map. In contrast, the basal connecting structures are much thicker and tilted at a sharp angle away from the radius of the axoneme, and these may well correspond to the “Y-linkers” frequently observed in conventional EM of cilia. Although the molecular composition of these structures is unknown, their location at the base of the CC suggests that proteins proposed to localize to that region are candidates. A number of the proteins we localized by STORM are seen in that basal region; however, none are restricted to or concentrated in that region.

Our three-dimensional map of the rod cilium obtained by cryo-ET provides a framework within which to place the molecular map revealed by STORM. Distinct radial compartments were identified by STORM, corresponding to the lumen of the axoneme, the axoneme itself, the space between the axoneme and the ciliary membrane, and finally, the membrane itself surrounded by the glycocalyx. While STORM reconstruction of centrin-2 signal clearly demarcates the lumen of the axoneme with a fairly uniform intensity along the length of the CC, the mechanism for this tightly restricted localization of this soluble centrin proteins is not known. In tomograms that have not been averaged, proteinaceous masses of irregular shape are found in this region and are connected by filaments to the walls of the axoneme, and these likely contain centrin-2 and other cilium-resident centrins. Acetylated tubulin labeling was similarly uniform along the length of the axoneme, but wider, consistent with the microtubule doublets of the axoneme surrounding the lumenal centrin-2. All other markers displayed much less uniform patterns along the axoneme, a feature not visible by lower resolution fluorescence techniques.

STORM reconstruction of BBS5 and STX3 to the CC provides unprecedented subciliary localization detail for these proteins in rod cilia. BBS5 is unique among BBS proteins as it features 2 PH-like domains that bind to phosphoinositides and is very likely the membrane-associated subunit of the BBSome (Jin et al., 2010). BBS5 and other BBS protein subunits of the BBSome were co-purified from OS preparations of retinal extract (Seo et al., 2011). Furthermore, BBS5 localization matches the STORM reconstruction of BBS9 at the CC (Figure S3), and thus BBS5 immunolabeling localizes the full BBSome complex in rod cells in our STORM reconstructions. Currently, the dearth of available BBS antibodies for immunofluorescence applications significantly hinders subcellular analyses of BBSome subunit dynamics in rod cilia.

In wild-type STORM reconstructions, BBS5 localizations cover the entire radius of the CC cytoplasm layer and extend into the membrane/WGA layer (Figure 2, S3). As the extended radius measurement of BBS5 accounts for the furthest diffuse single localizations from the CC lumen, the overall membrane localization of BBS5 is likely still within the cytoplasmic layer; however, BBS5 was previously localized along the periciliary membrane via immuno-EM (Smith et al., 2013), which is directly adjacent to the CC membrane. The contribution of the periciliary region to overall rod cell integrity has been demonstrated for Usher’s syndrome proteins: usherin, whirlin, and VLGR1 (Liu et al., 2007b; Yang et al., 2010), which are proposed to maintain a CC/periciliary interface (Maerker et al., 2008). IFT proteins are also localized to the periciliary region of the IS, where they putatively function in transport vesicle trafficking (Sedmak and Wolfrum, 2010). Thus, periciliary expression could contribute to STORM membrane localizations of BBS5. However, it is uncertain if the periciliary membrane maintains contact with the CC in all preparations, and, notably, these BBS5 STORM reconstructions do not favor strong localization to one face of the CC as would be predicted from the model of the periciliary/CC membrane interface.

The application of a compatible TUBA1 antibody for triple immunolabeling facilitated the analysis of BBS5 and IFT88 colocalization at the CC with STORM super resolution. BBS5 and IFT88 STORM reconstructions were consistently colocalized along the TUBA1 CC boundary with an outward orientation of BBS5 localizations surrounding the more centralized IFT88 clusters (Figure 3, S4). This colocalization was also apparent in the BBS and OS, as well. Together, these results are evidence for an *in vivo* interaction between the BBSome and the IFT complex as active transporters in the rod cilia. The BBSome has been hypothesized to be an “adaptor protein” for retrograde IFT complex transport based on trafficking defects in *Chlamydomonas* flagella (Baker et al., 2008; Lechtreck et al., 2009). In addition, the Arl6/BBS3 GTPase interacts with both the BBSome and IFT27, a Rab-like GTPase component of the IFT-B complex that is necessary for proper BBSome export from primary cilia (Liew et al., 2014). Furthermore, these interactions potentially link Arl6-mediated BBSome polymerization into a coat assembly and the incorporation of the BBSome with the IFT-B complex.

The lack of subciliary mislocalization of BBS5 at the CC in any of three BBS mutants analyzed (*Bbs2*^-/-^, *Bbs4*^-/-^ & *Bbs7*^-/-^ (Figure 6, S6) is particularly surprising, considering the significant morphological defects in rods associated with the BBSome (Datta et al., 2015; Gilliam et al., 2012). Loss of BBS1 or BBS7 severely disrupts BBSome formation *in vivo* (Seo et al., 2013; Zhang et al., 2013), but when BBS4 is deleted, the remaining BBSome subunits compensate to maintain a complex (Zhang et al., 2012). Therefore, the effect of losing individual BBS subunits on BBSome localization or function is unclear, especially considering that *Bbs* knockout mutations lead to dramatic morphological defects and rod cell death. Therefore, the effect of missing BBSome subunit proteins in mutant rod cilia, including the loss of “core” BBS2 and BBS7 subunits, is slight and leading to perhaps only gradual defects in CC trafficking which result in benign membrane accumulations that eventually become intolerable and trigger apoptosis. The less apparent OS accumulation of BBS5 compared to STX3 suggests that proper CC localization of the BBSome is not entirely BBSome-dependent. Or, rather, these results suggest that, functionally, the BBSome complex can at least partially compensate for missing subunits in the CC. Finally, it will be interesting in future analyses to consider if IFT protein CC localization is also maintained in *Bbs* mutant photoreceptors.

In STORM reconstructions of STX3 in WT rods, STX3 localization at the base of the OS confirms that this region is the likely destination for the t-SNARE protein (Figure 5, S3D). Based on a reported interaction with SARA near the basal disks of rat rod photoreceptors, STX3 has been proposed to be involved in disk formation and rhodopsin translocation into disks (Chuang et al., 2007), and in Xenopus, STX3 has been shown to have a key OS targeting motif in its SNARE homology domain (Baker et al., 2008). Besides rhodopsin (Wolfrum and Schmitt, 2000), no other clear localization data for proteins at the CC membrane has been described, and thus our STORM reconstruction of large clusters of STX3 at the CC membrane layer is novel localization data for any protein cargo trafficking along the CC. These large CC STX3 structural clusters may represent trafficking via mechanisms that are distinct from those involving IFT and the BBSome, whose markers, IFT81/88 and BBS5, have clearly distinct labelling patterns.

In all three *Bbs* knockout mutants used in this study, STORM subciliary localization of STX3 at the CC is attenuated in morphology but maintained in radial localization despite gross OS mislocalization (Figure 7 & S7). In these *Bbs* mutant rods, the predominant reconstruction of smaller morphological STX3 clusters might indicate either compensatory trafficking structures containing STX3 or the unidirectional anterograde STX3 trafficking only, as retrograde trafficking is very likely defective with *Bbs* knockouts. Notably, there are some smaller STX3 clusters along the CC axoneme in wild-type rods, which could be anterograde STX3 trafficking cargo, while the larger clusters may represent retrograde cargo (Figure 5, S3D). Another SNARE protein, STXBP1 (syntaxin binding protein 1), is also significantly mislocalized to the OS in *Lztf1/BBS17* and *BBS1*^*M390R*^ mutants, whereas Snap25 and VAMP are not mislocalized, despite also being tSNARE proteins expressed in rod cells (Datta et al., 2015). As such, the role of these proteins in the OS and their interaction with the BBSome remain subjects for future studies.

For future localization endeavors, nanoscopy methods continue to advance with technologies suited for studying the dynamics of the connecting cilium and subcellular photoreceptor biology. These include a Correlated Light and Electron Microscopy (CLEM) approach that aligns nanoscopy reconstructions onto EM micrographs for combined localization and morphological analysis (Johnson et al., 2015; Kim et al., 2015). Also, advances in 3D nanoscopy have achieved 3D super resolution through advanced imaging and reconstruction of serial ultrathin sections (Sigal et al., 2015), as well as through 4PI configured microscopes that reconstruct highly-accurate 3D localization maps (Huang et al., 2016). Our results establish the feasibility of combining nanoscopy with genetic models and other approaches toward analysis and observation of molecular events in mammalian rod cilia.

## MATERIALS AND METHODS

### Animals

All wild-type mice used for this study were C57BL/6 between ages 3 weeks and 3 months. For STORM immunostaining conditions, at least 3 WT mouse replicates were used; N-values in the text represent the number of individual rod cilia/connecting cilia analyzed per condition. *Bbs2*^-/-^ and *Bbs7*^-/-^mutant backcrossed C57BL/6 strains were acquired from the Jackson Laboratory (BBS2 – stock no. 010727, BBS7 – stock no. 24979) *BBS4*^-/-^ mice (also C57BL/6) were acquired from Dr. Samuel Wu and are described in (Eichers et al., 2006). Heterozygous crosses were bred to produce homozygous knockouts from each strain as determined by mouse tail genotyping. Multiple age timepoints were tested for STORM mutant analyses of *Bbs4*^-/-^ mice (5 weeks, n=4; 8 weeks n=4) and *Bbs7*^-/-^ mice (7 weeks, n=2; 8 weeks, n=4). *Bbs2*^-/-^ mice were only used at the 8 week timepoint (n=3) as there was a lower rate of viable knockouts from this strain in our colony. All procedures were approved by the Baylor College of Medicine Institutional Animal Care and Use Committee.

### Cryo-ET

Rod cell fragments containing outer segments, CC, and portions of the IS were collected from WT mice by iso-osmotic density-gradient centrifugation as described previously (Gilliam et al., 2012; Wensel and Gilliam, 2015). A suspension of cells in Ringer’s buffer (20 μL) was mixed with 8 μL of BSA-stabilized 15 nm fiducial gold (Electron Microscopy Sciences), and 2.5 μL of the mixture was applied to a pre-cleaned, glow-discharged 200 mesh Quantifoil carbon-coated holey grids which had been pre-cleaned and glow discharged. Samples were plunge-frozen in liquid ethane held at liquid nitrogen temperature using a Vitrobot Mark III automated plunge-freezing device. Samples were imaged in a JEM2100 electron microscope with a Gatan cryo-stage at −180 °C. The frozen-hydrated specimen were imaged at −180 °C in a JEM2100 electron microscope (Gatan Company), equipped with a 4k x 4k CCD camera, and a Polara G2 electron microscope (FEI Company), equipped with a field emission gun and a direct detection device (Gatan K2 Summit). Using the JEM2100 electron microscope, images were collected at magnifications of 12,000 x at 200 kV with the defocus of ~7-10μm. Serial EM was used to collect single-axis tilt series of +/- 60° with 2° increments. The cumulative electron dose was 50-80 e^−^/Å^2^. Micrograph alignment and 3D reconstruction were performed using IMOD (Mastronarde, 1997; Mastronarde and Held, 2017). Using the Polara G2 electron microscope, images were collected at magnifications of 12,000 x at 300 kV with a defocus of ~9 μm. Single-axis tilt series with a cumulative dose of ~50 e^−^/å^2^ distributed over 35 stacks covering an angular range of −51° to +51° with 3° increments. The drift correction of the dose-fractionated tilt series was corrected using Motioncorr. Alignment and 3D reconstruction were again performed using IMOD (Mastronarde, 1997; Mastronarde and Held, 2017). Subtomogram averaging was carried out using custom software following procedures described previously (Koyfman et al., 2011).

### Antibodies and Labeling Reagents

The BBS5 monoclonal antibody used in this work is described in (Smith et al., 2013) with validation. Commercial primary antibodies included in this study were: anti-acetylated alpha-tubulin (mo MC, clone 6-11B-1, Santa Cruz, sc-23950), anti-centrin-2 (rab PC, sc-27793), anti-centrin-2 (rab PC, Proteintech, 15866-1-AP), anti-Arl13b (rab PC, Proteintech, 17711-1-AP), anti-IFT88 (rab PC, Proteintech, 13967-1-AP), anti-IFT81 (rab PC, Proteintech, 11744-1-AP), anti-BBS9 (rab PC, sc-292152), anti-TUBA1-ATTO488 (human MC, antibodies-online.com, ABIN4888971), anti-STX3 (rab PC, Proteintech, 1556-1-AP). 10 μg of Wheat Germ Agglutinin conjugated to Alexa 647 (Molecular Probes) was used as a secondary antibody in immunostaining procedures for glycoprotein labeling of mouse retinas.

### Confocal Immunohistochemistry and Imaging

All antibodies and labeling reagents used for immunofluorescence were verified for positive signal with confocal microscopy (Figure S4E). Cryofixation of mouse eyes for connecting cilia immunostaining is described in (Bolch et al., 2016). Mouse eye cups are embedded in Optimal Cutting Temperature (OCT) media and plunged into freezing isopentane (i.e. 2-Methlybutane, Sigma) to cryo-fix for 30 minutes before cryosectioning at 8 μm thickness. Sections were blocked in 2% Normal Goat Serum (NGS, Fitzgerald Industries) + 2% Fish Scale Gelatin (Sigma) + 2% Bovine Serum Albumin (Sigma) + 0.2% Triton X-100 dilute in 1x PBS and probed with 0.5 - 1 μg of primary antibody at room temperature overnight. After secondary antibody labeling with goat anti-mouse IgG-Alexa 488 (Thermo Fisher) and/or goat anti-rabbit IgG-Alexa 555 (Thermo Fisher) (1:500), sections were mounted with Vectashield (Vector Laboratories) for imaging on a Leica TCS-SP5 confocal microscope.

### STORM Immunohistochemistry and Resin Embedding

Retinas from either WT or *Bbs* mutant mice were immunolabeled for STORM in a modification of whole mount staining, in which whole retina were stained in solution following a two-step protocol. First, retinas were dissected unfixed in ice cold Ames’ media (Sigma) and immediately blocked in 10% NGS (Fitzgerald Industries) + 0.3% saponin (Sigma) + 1x Protease Inhibitor Cocktail (GenDepot) diluted in 1x Ames’ media for 2 hours at 4°C. Unfixed labeling of the retina is necessary for complete antibody penetration of the connecting cilium axoneme, which has been described previously (Evans et al., 2010; Hidalgo-de-Quintana et al., 2015). Primary antibodies (5-10 μg) were added to the blocking buffer and incubated at 4°C for 20-22 hours. Retinas were washed 3 times for 5 minutes in 2% NGS in Ames’ media on ice before secondary antibodies were added to the same buffer and incubated at 4°C for 2 hours. Secondaries used (8 μg each): F(ab’)2-goat anti-mouse IgG Alexa 647 & F(ab’)2-goat anti-rabbit IgG Alexa 555 (Thermo Fisher). In WGA-Alexa647 labeling experiments, F(ab’)2-goat anti-rabbit IgG Alexa 555 (Thermo Fisher) was used for dual labeling. Retinas were washed in 2% NGS/Ames 6 times for 5 minutes each on ice and fixed in 4% formaldehyde diluted in 1xPBS for 15 minutes at room temperature.

Next, retinas were re-blocked in 10% normal goat serum + 0.2% Triton X-100 diluted in 1xPBS for 2 hours at room temperature. Primary antibodies (5-10 μg each) were re-added to the blocking buffer and incubated for 2 days at 4°C. This second IHC step features Triton permeabilization to assure complete antigen binding in and around the connecting cilium. After the second primary antibody incubation, retinas were washed 4x for 10 minutes each in 2% NGS/1x PBS. The same secondary antibodies were added to the wash buffer as before (8 μg each) for overnight incubation at 4°C. Retinas were washed 6x in 2% NGS/1x PBS for 5 minutes each before postfixation in 3% formaldehyde diluted in 1x PBS for 1 hour at room temperature.

The resin embedding protocol for immunolabeled retinas is based on (Johnson et al., 2015; Kim et al., 2015; Sigal et al., 2015). Post-fixed retinas were dehydrated in a series of ethanol washes (15 minutes each: 50%, 70%, 90%, 100%, 100%) followed by embedding steps of increasing concentrations with Ultra Bed Epoxy Resin (Electron Microscopy Sciences) to ethanol (2 hours each: 25%:75%, 50%:50%, 75%:25%, 100% resin twice). Embedded retinas were cured on the top shelf of a 65’C baking oven for 20 hours. 500 nm - 1 micron sections were cut on a UCT or UC6 Leica Ultramicrotome and dried directly onto glass-bottom dishes (MatTek 35 mm dish, No. 1.5 coverslip)

### STORM Image Acquisition

Immediately prior to imaging, 10% sodium hydroxide (w/v) was mixed with pure 200-proof ethanol for 30 minutes to prepare a mild sodium ethoxide solution. Glass-bottom dishes with ultra-thin retina sections were immersed for 30-40 minutes for chemical etching of the resin that facilitates STORM in embedded tissue (Johnson et al., 2015; Kim et al., 2015; Sigal et al., 2015). Etched sections were then washed and dried on a 50°C heat block. The following STORM imaging buffer was prepared: 50 mM Tris (pH 8.0), 10 mM NaCl, oxygen scavenging system: 0.56 mg·ml^−1^ glucose oxidase (Amiresco) + 34 μg ml-1 catalase (Sigma), 10% (w/v) glucose + 15 mM MEA (i.e. L-cysteamine, Chem-Impex) + 10% Vecta Shield (Vector Laboratories). Imaging buffer was added onto the dried, etched sections and sealed with a second number 1.5 coverslip for imaging.

Imaging was performed on the Nikon N-STORM system, which features a CFI Apo TIRF 100x oil objective (NA1.49) on an inverted Nikon Ti Eclipse microscope that houses a quad cube filter (Chroma, zt405/488/561/640 m-TRF) and a piezo Z stage with Nikon’s Perfect Focus System for Z stability. An Agilent MLC400B laser combiner with AOTF modulation housed the 200 mW 561 nm and 647 nm solid-state lasers that were used for imaging in this study. STORM image acquisition was controlled by NIS-Elements Ar software. For each STORM acquisition, a 512×512 pixel field was captured by an Andor iXON DU 897 EMCCD camera (pixel size = 160 nm wide for roughly a 40 square micron STORM area), and a cylindrical lens was inserted into the light path (see below). Chromatic aberration between channels was corrected via X-Y warp calibration.

The etched thin sections were first scanned with low laser power to locate a region with multiple and sufficiently bright CC. From this region, DIC images and low power laser images (widefield fluorescence images) were saved for reference. Both the 561 nm and 647 nm laser lines were increased to maximum power to initiate photoswitching. Using an independent power meter, maximum laser power was measured directly above the objective as 34.8 mW for 561 nm and 65.5 mW for 647 nm. Imaging frames were collected at ~56 frames per second. 20,000 - 50,000 frames were collected for each imaging experiment. 561 nm & 647 nm frames were collected sequentially without any lower wavelength “activation” light (a STORM protocol previously termed direct-STORM or dSTORM)

### STORM Image Analysis

2D-STORM Analysis of STORM acquisition frames was performed using NIS Elements Ar Analysis software with drift correction. Analysis identification settings were used for detection of the individual point spread function (PSF) of photoswitching events in frames from both channels to be accepted and reconstructed as 2D Gaussian data points. These settings were as follows: Minimum PSF height: 400, Maximum PSF height: 65,636, Minimum PSF Width: 200 nm, Maximum PSF Width: 700 nm, Initial Fit Width: 350 nm, Max Axial Ratio: 2.5, Max Displacement: 1 pixel. These screening parameters correspond to ellipsoid Gaussian fitting, which yielded stronger reconstructions from ellipsoid PSF’s generated by the cylindrical lens is inserted into the light path compared to circular Gaussian fitting and no cylindrical lens. Notably, fluorescence within the entire CC structure (300 - 400 nm) is within the effective depth of field of the microscope and thus 2D-STORM reconstructions are effectively super resolution 2D projections of entire connecting cilia. 3D-STORM via astigmatism was not successful in our samples due to background inherent to our tissue preparation, which rendered poor Z localization.

After analysis, reconstruction quality was assessed by plotting Gaussian localization accuracy based on the Thompson equation (Thompson et al., 2002). For each reconstructed connecting cilia Gaussian cluster (defined as a selected ROI surrounding a connecting cilium and surrounding structures), minimum photon filter values were increased or decreased and a local density filter (typically set to 20 molecule thresholding value at a 50-100 nm radius) was used to eliminate background localizations to attain an average localization accuracy per molecule of <20 nm in each reconstruction, which was used a benchmark for good STORM reconstruction and acceptable background. Reconstructions were processed in Fiji/ImageJ, and the Straighten tool was applied to straighten curved or bent cilia to acquire accurate length profiles. Look up table (LUT) settings for visualization within the analysis software were removed from all reconstructions to limit the saturation of strong Gaussians; however, whole image contrast was adjusted for clarity when necessary.

In Fiji/ImageJ, ROI’s of digitally straightened STORM reconstructions were measured using row average profiling, which plots the average intensity across the width of the ROI for each row of pixels along the length of the ROI. Pixels were converted to nm for accurate scaling. From these row average profiles, the edges of STORM clusters were set as 1/e times the maximum intensity value for any given cluster profile, and distance from center (radius) was measured as the max intensity value for either acTub and centrin-2 CC clusters (the core) to the edge (1/e max) of the STORM cluster of interest. All presented profiles are normalized by area under the curve. All measurements were made in a 1.1 μm longitudinal region just above the basal body that corresponds to the length of the ultrastructural CC and provided as mean ± standard deviation. For distance from center measurements, STORM clusters that extended beyond either the acTub or centrin-2 cluster were measured instead of more colocalized clusters. All measurements were rounded to the nearest nanometer to account for several contingent factors that negate sub-nanometer accuracy of reconstructed STORM clusters, including antibody displacement (antibody “linkage error”). Based on our measurements, the reconstructed connecting cilia are slightly widened based again on linkage error, or other factors such as waviness of individual cilia, slight flattening of the structure by the embedding media and convolution with the point-spread function at the edges of measured STORM clusters.

## ACKNOWLEDGMENTS

This work was supported by NIH grants R01-EY007981, R01-EY026545, and the Welch Foundation Q0035 to TGW, by the National Center for Macromolecular Imaging, P41-GM103832 (MFS), by NIH fellowships F31-EY028025 (VLP) and F32-EY027171 (MAR), and by the BCM Vision Research Core P30-EY002520. The authors thank Dr. Ivan Anastassov, Dr. Melina Agosto and Mr. Christopher Hampton for useful insight and discussion. The authors also thank Dr. Clay Smith for sharing resources and Mr. Matthew Mitschelen (Nikon) for technical advice and guidance.

